# Prevalence of underweight and it’s associated factors among children aged 6-59 months in Angolalla Tera district, Northeast Ethiopia

**DOI:** 10.1101/2020.07.23.217349

**Authors:** Leweyehu Alemaw Mengiste, Yoseph worku, Endeshaw Degie Abebe, Wondimeneh Shibabaw shiferaw

## Abstract

**Introduction:** Undernutrition is a major public health problem all over the world. In Ethiopia, the child malnutrition rate is one of the most serious public health problems and the highest in the world. High malnutrition rates in the country pose a significant obstacle to achieving better child health outcomes.

**Objectives:** This study was aimed to assess the prevalence of underweight and its associated factors among 6-59months of age children in Angolela Tera district, northeast Ethiopia.

**Methods:** A community based cross-sectional study was conducted by a simple random sampling technique with a sample size of 414 enrolled mothers with 6-59months of children. Semi-structured questionnaires were used to collect data. The data was entered using EPI DATA version 3.1and analysis was done by SPSS version 24 and WHO Anthro software was used for anthropometry calculation. Bivariable and multivariable logistic regression analysis was used.

**Result:** Among 414 mothers with child pairs the result of the current study indicated that 15.9% (95% CI: 12.6-19.6)were underweight. Being male (AOR=1.8;95%CI;1.04-3.16), birth interval below 24 months (AOR3.2=95%CI;1.59-6.71), average monthly family income less than 1596ETB (AOR4.9=95%: CI;2.53-9.76), Children having diarrhea in the past two weeks before the data collection (AOR =9.06; 95% CI: 3.14-26.12), and children having diarrhea within two weeks (AOR=2.06;95%: CI;1.07-3.96) were significantly associated with underweight.

**Conclusion:** This study revealed a high prevalence of underweight among children aged 6-59 months in the study area. All the concerned bodies should be strengthening the health extension program to improve and provide the necessary education for the community on nutritional programs, environmental sanitation, and diarrhea prevention.

## Background

Malnutrition is one of the major public health problems all over the world. Currently, it faces and associated with more than 41%of the deaths that occur annually in children from 6 to 24 months of age in developing countries, which were approximately 2.3million (1). Globally, undernutrition contributes more than one-third of child deaths which can be prevented through public health interventions(2, 3).

Undernutrition can affect children’s health and learning ability during their adulthood life. Children with undernutrition are usually suffered from chronic illnesses(4). Survival of malnutrition can suffer from impaired physical development and intellectual abilities, which in turn may diminish their working capacity with negative effects on economic growth. Child malnutrition may also lead to higher levels of chronic illness in adult life and these may have intergenerational effects, as malnourished females are more likely to give birth to low-weight babies(5).

Underweight, defined based on weight-for-age, is a composite measure of stunting and wasting and is recognized as the indicator for assessing changes in the magnitude of malnutrition over time. (WHO) Underweight remains one of the most common causes of morbidity and mortality among children throughout the world(6, 7). Malnutrition is one of the leading causes of morbidity and mortality in children under the age of 5 years in developing countries. Every year, 3.5 million children died due to malnutrition-related causes, of which underweight accounts for nearly 1 million (8, 9). About two in five (38%) children in sub-Saharan Africa are underweight; 10.5% wasted, (2.2% severely), and 46.5% stunted (half of them severely) In Ethiopia, several studies have demonstrated that the prevalence of underweight was 28.4% in the Amhara region, (11), 14.3% in the bure district (12).

Though childhood underweight continues to be the leading public health problem in developing countries, it can occur as a result of a wide range of factors. Various reports have been indicated that underweight in children is mainly caused by inadequate food intake (13, 14), repeated infections(15, 16), poor feeding practices (16), lack of ANC follow up (17, 18), rural residence (19), child-rearing practices (20), low economic status (20, 21), social, and cultural factors(22) In Ethiopia, the magnitude of underweight was still high. Evidences claimed that according to the 2016 EDHS report underweight was 41% in 2000, 33% in 2005, and 29% in 2011, and24 in 2016(23).

Knowledge of the prevalence of underweight and its contributing factors is an important prerequisite for developing strategies of nutritional intervention. Though, there is no recent study which identifies the prevalence and predictors of undernutrition in the study area. Therefore this study was designed to estimate the prevalence of underweight and its associated factors among children aged 6–59 months in Angolalla tera district, northeast Ethiopia

## METHODS

### Study design, setting and period

A community-based cross-sectional study was conducted from December 30 to January 30, 2018 The study was conducted in Angolalla Tera woreda North Shewa zone. This woreda is 112Km far from Addis Ababa the capital city of Ethiopia. The woreda has 21 kebeles, with a total area of 1,508.19Km3 hectare and the total population was 98,382 of these males comprise 50,458 and 47,924 females. Out of the 15,175 were under-five children, out of these about 7, 401 were males and 7, 774 were female. In the woreda, there is four health center and 21 health post. The livelihood of most of the population is earned from farming and employment from government and non-government organizations.

### Sample Size Determination and Sampling Techniques

The sample size was determined using a single population proportion formula assuming that 51.1% was the prevalence of stunting under-five children which were done in the Amhara region, Lalibela town northern Ethiopia in 2014(25). And a 5% margin of error with a 95% confidence level with anticipated a 10% non-response rate.

The required sample size was 384 and with adjustment for non-response rate (10%) the final required sample size was 422 mother-child pairs.

From 21 kebeles, six kebeles were selected by using simple random sampling techniques. Study participants/households/ were allocated to selected kebeles by proportionate allocation and from each selected kebeles. A simple random sampling technique was used to select children from households (table of random numbers) based on frame existing in health posts

### Populations

The source populations were all mothers with children pair from 6-59 months of age who live in the households of Angolalla tera district North Shoa, Amhara region. The study populations were all randomly selected children 6-59 months of age who were living with their mothers in the sampled kebeles of Angolalla tera district administration during the study period.

### Eligibility criteria

#### Inclusion criteria

Children 6-59 months of age who were living with their mothers and whose mothers were available in the selected households.

#### Exclusion criteria

Children, Child’s mother critically ill, during data collection who were seriously ill, had physical deformities of limbs and spines were excluded because of difficulty in height measurement.

## Study Variables

### Dependent variable

underweight status in children 6–59 months of age

### Independent variables

#### Socio-demographic factor

Age of child, Sex, marital status of the mother, birth order of the child, preceding birth interval of child, mother’s religion, family income and educational status

#### Environmental factors

Source of drinking water and availability of Latrine facility

#### Dietary factors

Ever breastfeeding, time for initiation of breastfeeding, Colostrum feeding, pre-lacteal feeding, duration of breastfeeding, age for introduction of complementary food, and method of feeding.

#### Health care factors

Child’s immunization status, child’s diarrhea, mothers antenatal care visits, mother’s age of pregnancy, and mother’s place to deliver

### Operational Definitions

#### Anthropometry

Measurement of the variation of physical dimensions and the gross composition of the human body at different age levels and degrees of nutrition by weight-for-age, height-for-age and weight-for-height(24).

#### Complimentary food

Foods that are required by the child, after six months of age, in addition to sustained breastfeeding.

#### Diarrhea

Diarrhea is defined for a child having three or more loose or watery stools per day.

#### Underweight

is defined as underweight if WAZ below the −2 SD from the NCHS/WHO reference of the median of the standard curve. A severely underweight was diagnosed if it was below −3 SD (24).

#### Pre-lacteal feeding

Newborns exposed to any foods, substances or drinks other than human milk before the initiation of breastfeeding or during the first three days of birth

#### Currently on vaccination

children who have received a vaccination according to the EPI schedule based on their age.

### Data collection tools and procedure

Data was collected by using a semi-structured questionnaire that was adapted from the United Nations Children’s Fund and previous similar literature. Questionnaires were prepared in English and translate into Amharic local language and then back to English to keep its consistency. The tool contains underweight, socio-demographic, environmental, healthcare, and dietary factors among children age (6-59months). Four diploma nurses were collect the data and two BSc nurses were take part in the supervision. The data collectors and supervisors were selected based on experience on research, familiarity with the study area, local language, and interest to participate in the study.

#### Anthropometric data

The anthropometric data were collected using the procedure stipulated by the WHO (2006) for taking anthropometric measurements. Before taking anthropometric data for children; their age should first determined to ensure the study population.

#### Age

The child’s age was interviewed from the mother and confirmed by using a birth certificate or vaccination cards and also we were used a “local-events” calendar

#### Weight measurement

Weight was measured by an electronic digital weight scale with minimum/lightly/clothing and no shoes. Calibration was done before weighing every child by setting it to zero. In the case of children age below two years, the scale was allowed weighing of very young children through an automatic mother-child adjustment that used to eliminate the mother’s weight while she standing on the scale with her baby (17).

### Data quality control, Data processing and analysis

Pre-testing was conducted on 5% of sample size at Basona Werana district before the actual data collection process. Training was given to the data collector and supervisor by the principal investigator for one day about the objectives of the study, data collection instruments, data collection procedures, physical measurement, and the ethical consideration during data collection.

The scales indicators were checked against zero reading after and before measuring the weight of the child. Daily collected information was reviewed and possible errors should be returned to the data collectors for correction. The investigator was supervised and reviews every questionnaire for completeness. Data entry, coding, and cleaning were performed by the principal investigator.

The collected data were checked for completeness, coded, and then entered to Epi Data Version 3.1 then exported to SPSS version 24 for analysis. Descriptive statistics were used to describe the study participants about relevant variables. To determine the actual predictors of underweight, binary logistic regressions were applied and the variables found to have a p-value of <0.2 with the outcome variable at bivariable analysis were exported to multivariate analysis which uses to control confounding effect. Moreover, the variables which have significant association will be identified based on p-values< 0.05 and AOR (adjusted odds ratio), with 95% CI to measure the strength of the associations. Assumption tests of logistic regression were held and the model goodness of fit of the multivariate logistic regression checked by using Hosmer and Lemeshow test.

## RESULT

### Socio-demographic characteristics of the study participants

A total of 414 respondents participated in yielding a response rate of 98.1%. The mean age±1 Standard Deviation of the participants was 24.9 (±15.6) months. Within this ample, 212 (51.2%) were males; about one-third 142 (34.3%) of participants, were with 12-23 months old age. The majority,87.4% of the participants were Orthodox Christians, First birth order children comprised 114 (27.7%) of the participants. The majority of the mothers were married 87.4%, housewives 87.7%, and nearly two thirds 62.3% were illiterate (Table 1)

**Table 1.**
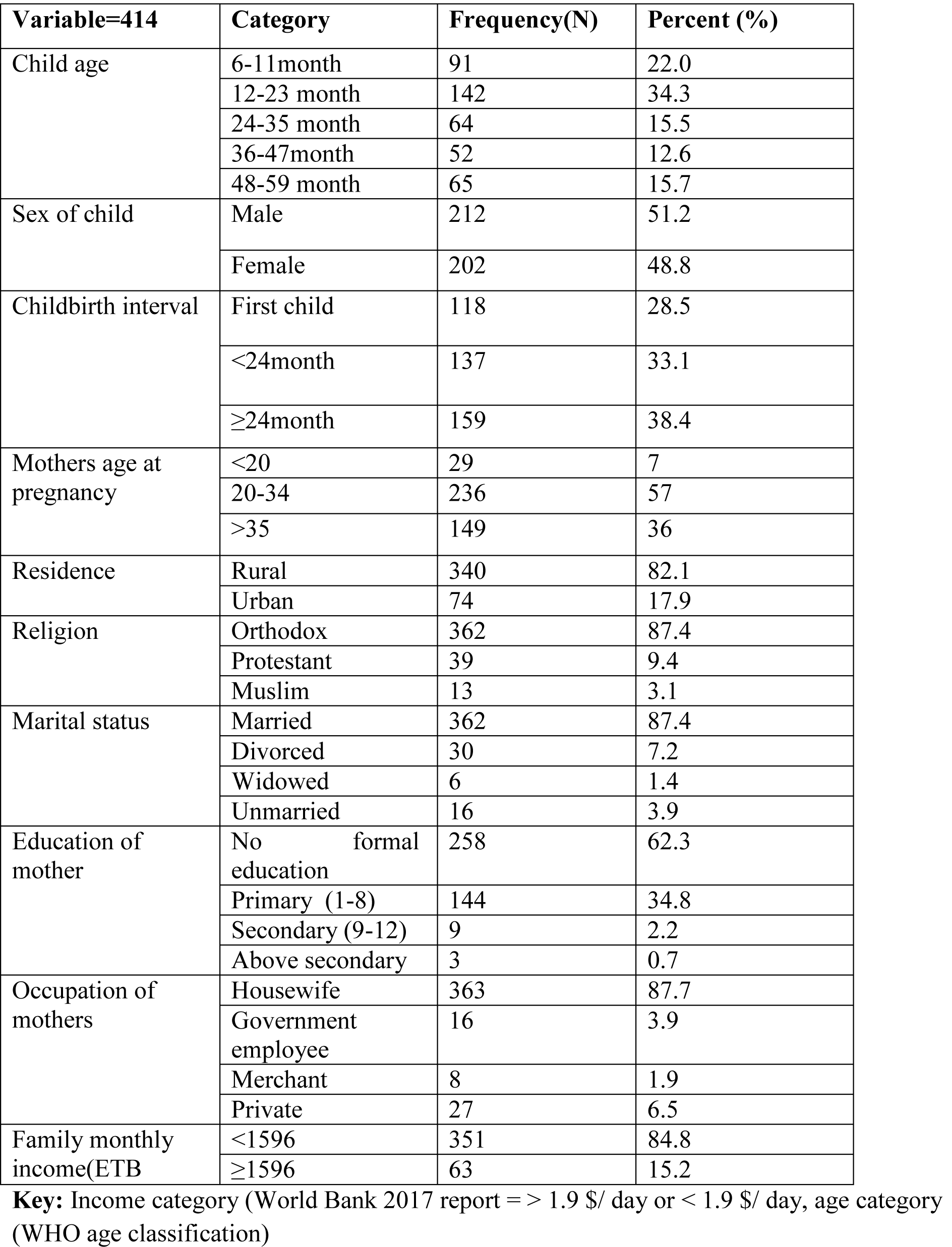
Socio-demographic characteristics of children among 6 to 59 months in Angolela Tera District, northeast Ethiopia, 2019 (N=414)

| **Variable=414** | **Category** | **Frequency(N)** | **Percent (%)** |
| --- | --- | --- | --- |
| Child age | 6-11month | 91 | 22.0 |
|  | 12-23 month | 142 | 34.3 |
|  | 24-35 month | 64 | 15.5 |
|  | 36-47month | 52 | 12.6 |
|  | 48-59 month | 65 | 15.7 |
| Sex of child | Male | 212 | 51.2 |
|  | Female | 202 | 48.8 |
| Childbirth interval | First child | 118 | 28.5 |
|  | <24month | 137 | 33.1 |
|  | ≥24month | 159 | 38.4 |
| Mothers age at pregnancy | <20 | 29 | 7 |
|  | 20-34 | 236 | 57 |
|  | >35 | 149 | 36 |
| Residence | Rural | 340 | 82.1 |
|  | Urban | 74 | 17.9 |
| Religion | Orthodox | 362 | 87.4 |
|  | Protestant | 39 | 9.4 |
|  | Muslim | 13 | 3.1 |
| Marital status | Married | 362 | 87.4 |
|  | Divorced | 30 | 7.2 |
|  | Widowed | 6 | 1.4 |
|  | Unmarried | 16 | 3.9 |
| Education of mother | No formal education | 258 | 62.3 |
|  | Primary (1-8) | 144 | 34.8 |
|  | Secondary (9-12) | 9 | 2.2 |
|  | Above secondary | 3 | 0.7 |
| Occupation of mothers | Housewife | 363 | 87.7 |
|  | Government employee | 16 | 3.9 |
|  | Merchant | 8 | 1.9 |
|  | Private | 27 | 6.5 |
| Family monthly income(ETB | <1596 | 351 | 84.8 |
|  | ≥1596 | 63 | 15.2 |

**Key:** Income category (World Bank 2017 report = > 1.9 \$/ day or < 1.9 \$/ day, age category (WHO age classification)

### Error! Reference source not found.Health care and environmental characteristics

About, 185(44.7%) of children had normal birth weight (2.5-4.0 Kg) and 50(12.1%) were <2.5Kg. The majority, 395 (95.4%) of children were immunized; out of these 183(46.3%) were fully vaccinated. More than two-third, 288(69.6%) of the participant had got diarrhea during two weeks of the period before data collection. More than half,233(56.3%) of the mothers had no antenatal care visits. Nearly half, 178(43%) of the households used protected well as the main source of drinking water, (Table 2).

**TABLE 2.**
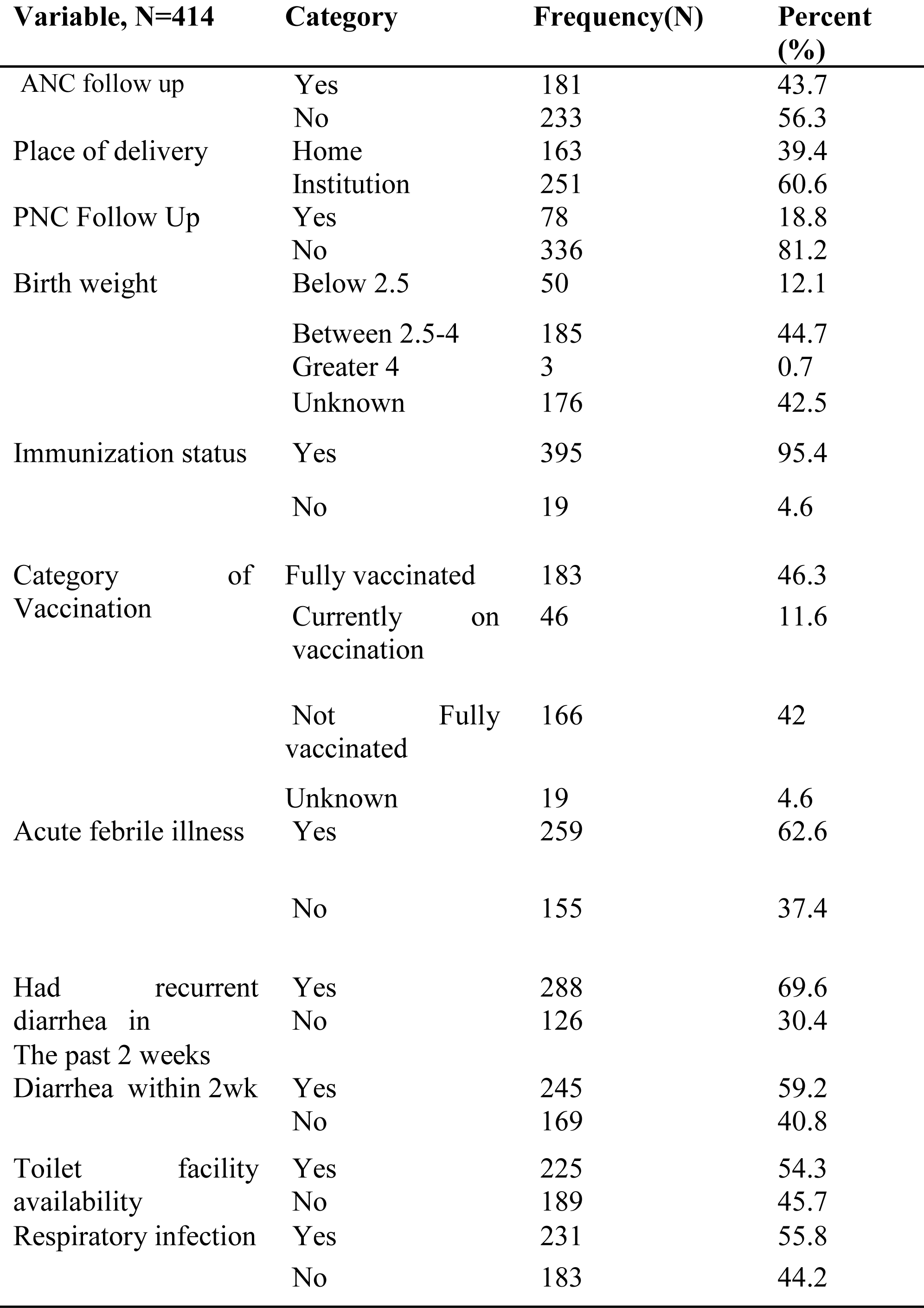
Environmental and health care characteristics of children.

### Dietary characteristics

Regarding breastfeeding practice majority 380 (91.8%), of children were breastfed in the first six months; nearly half 199(46.1%) of children started breastfeeding within the first one hour of birth. Most of the study participants 314(75.8%) were receive colostrum. Children who breastfeed for less than 12 months were 191(46.1%), 13 −24 months 167 (40.3%), and more than 12 months were 56(13.5%). About two-thirds of respondents 281(67.9%) started complementary feeding at the age of 6 months. Concerning the method of feeding, mothers who used a cup to feed their children were 140 (33.8%), and 138 (33.3%) used a hand to feed their children, (Table3).

**TABLE 3;.**
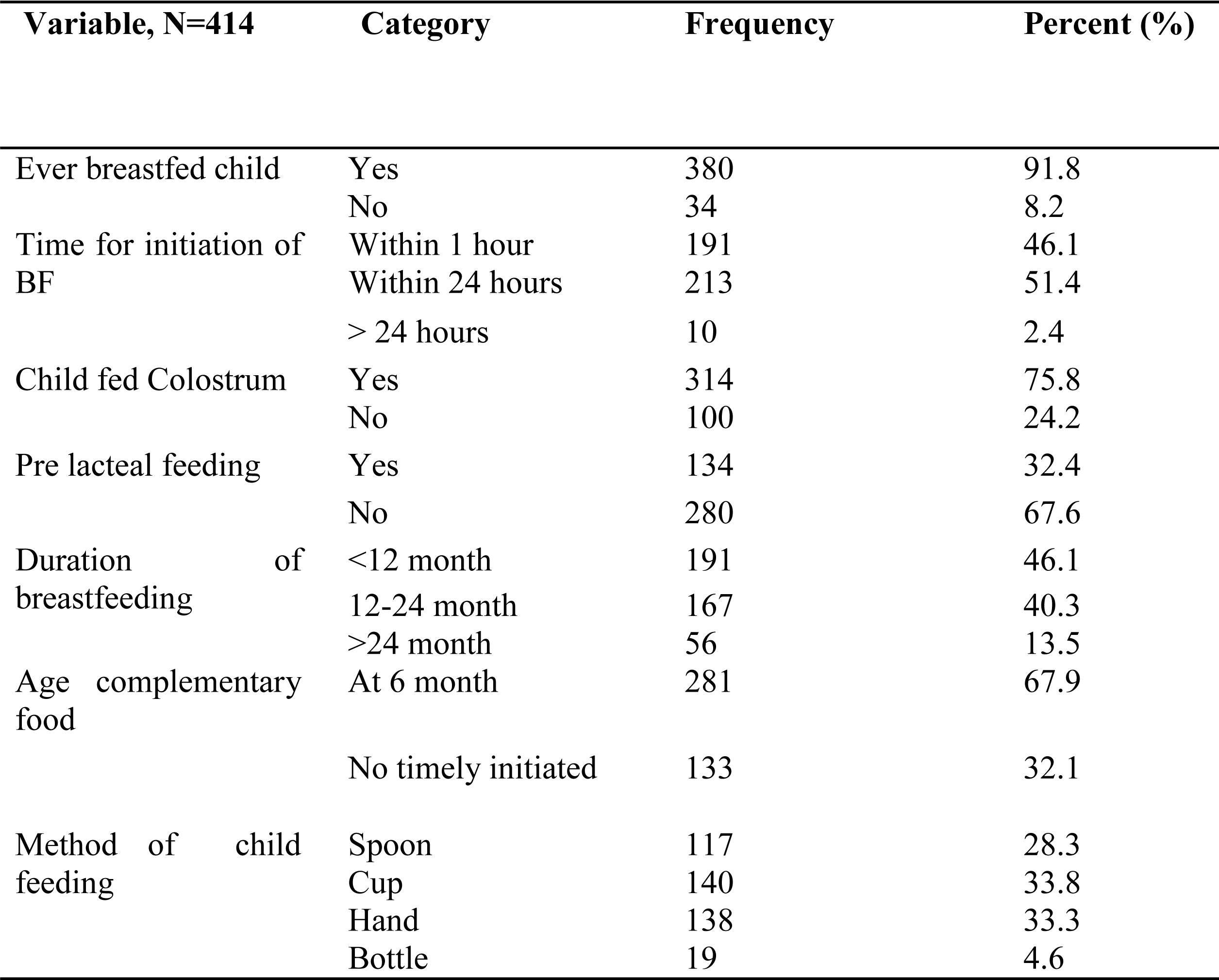
Dietary characteristics of children.

| Variable, N=414 | Category | Frequency | Percent (%) |
| --- | --- | --- | --- |
| Ever breastfed child | Yes | 380 | 91.8 |
|  | No | 34 | 8.2 |
| Time for initiation of BF | Within 1 hour | 191 | 46.1 |
|  | Within 24 hours | 213 | 51.4 |
|  | > 24 hours | 10 | 2.4 |
| Child fed Colostrum | Yes | 314 | 75.8 |
|  | No | 100 | 24.2 |
| Pre lacteal feeding | Yes | 134 | 32.4 |
|  | No | 280 | 67.6 |
| Duration of breastfeeding | <12 month | 191 | 46.1 |
|  | 12-24 month | 167 | 40.3 |
|  | >24 month | 56 | 13.5 |
| Age complementary food | At 6 month | 281 | 67.9 |
|  | No timely initiated | 133 | 32.1 |
| Method of child feeding | Spoon | 117 | 28.3 |
|  | Cup | 140 | 33.8 |
|  | Hand | 138 | 33.3 |
|  | Bottle | 19 | 4.6 |

### Prevalence of underweight among children

In this study, the prevalence of underweight among children age 6-59 months was 15.9%. also, our findings showed that the prevalence of severe underweight among the children was 25(6.0%) Figure1, furthermore, the prevalence of underweight was peaking at 12-23 months, which was 20(30.3%), (Figure 2).

**Figure 1.**
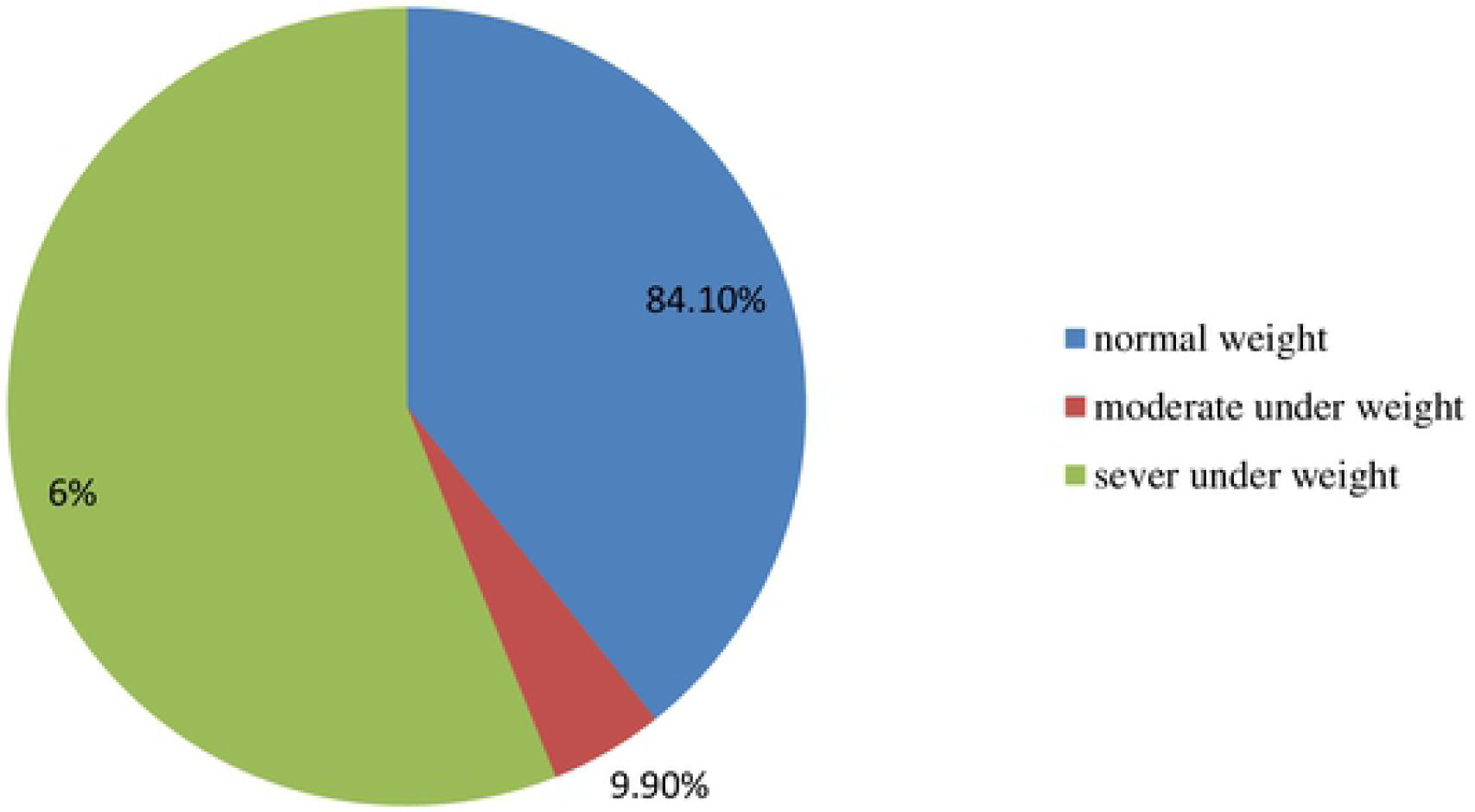
prevalence of underweight Among six to fifty-nine months of age in Angolela Tera district, North Shoa, Amhara, Ethiopia, 2019 (N=414)

### Associated factors of underweight

Based on the multivariable logistic regression analysis of this study, being male, birth interval below 24 months, family income less than 1596ETB children having diarrhea within two weeks, and diarrhea incidence before two weeks of data collection were significantly associated with underweight. Male children were 1.81 times (**AOR=1**.**81; 95% CI: 1**.**04-3**.**16**) more likely to be underweight than females children. =on the other hand, children those family average monthly income less than1596 birr were 4.9 times (**AOR=4**.**97; 95%CI: 2**.**53-9**.**76**) more likely to be underweight than those family income greater than1596ETB.Regarding the association of birth interval with underweight, mothers whose birth below 24 months were 3.2times more likely to be underweight than those birth space more than 24 months (**AOR=3**.**27; =95%CI; 1**.**59-6**.**71**).Children having diarrhea in the past two weeks before the data collection were 9 times (**AOR =9**.**06; 95% CI: 3**.**14-26**.**12**) more likely to develop underweight than children without diarrheal disease. Children having diarrhea within two weeks before the data collection were 2.06 times (**AOR =2**.**06, 95% CI: 1**.**07-3**.**96**) more likely to develop underweight than children without diarrheal disease (**Table 4)**.

**Table 4:** Bivariable and multivariable logistic regression analysis of factors associated with underweight.

| Variable | Underweight<br>Yes | No | COR(95%CI) | AOR(95%CI) |
| --- | --- | --- | --- | --- |
| Sex |  |  |  |  |
| Male | 42(63.6) | 170(48.9) | 1.83(1.06-3.156) | 1.81(1.04-3.16)* |
| Female | 24(36.4) | 178(51.1) | 1 | 1 |
| Residence |  |  |  |  |
| Rural | 58(87.9) | 282(81.0) | 1.69(0.77-3.72) | 1.64(0.72-3.71) |
| Urban | 8(12.1) | 66(19.0) | 1 | 1 |
| Birth interval |  |  |  |  |
| 1first child | 18(27.3) | 100(28.7) | 2.02(0.94-4.31) | 1.68(0.75-3.74) |
| <24month | 35(53.0) | 102(29.3) | 3.85(1.94-7.72) | 3.27(1.59-6.71)* |
| ≥24 month | 13(19.7) | 146(42.0) | 1 | 1 |
| Income |  |  |  |  |
| <1596 | 44(66.7) | 307(88.2) | 3.74(2.04-6.86)) | 4.97(2.53-9.76)* |
| ≥1595ETB | 22(33.3) | 41(11.8) | 1 | 1 |
| Ill Diarra<br>within two<br>week |  |  |  |  |
| Yes | 53(80.3) | 192(55.2) | 3.31(1.74-6.29) | 2.06(1.07-3.96)* |
| No | 13(19.7) | 156(44.8) | 1 | 1 |
| Diarra before<br>2week |  |  |  |  |
| Yes | 62(93.9) | 226(64.9) | 8.36(2.97-23.55) | 9.06(3.14-26.12)* |
| No | 4(6.1) | 122(35.1) | 1 | 1 |
| ANC |  |  |  |  |
| YES | 19(28.8) | 162(46.6) | 0.46(0.26-0.83) | 0.554(0.31-1.01) |
| NO | 47(71.2) | 186(53.4) | 1 | 1 |

## Discussion

This study revealed a high prevalence of child underweight in the study area. The finding showed that 15.9 %(95% CI: 12.6-19.6) of children were underweight. This finding is in line with studies conducted on Beta-Israel children in the Amhara region, 14.6% o (26), and 15.8% in Hawasa (4). However, our finding is lower than the 2016 EDHS report in which 24% of the children underweight (28), 27% in Nepal (29), 27.7% in Vietnam (30).

2016 EDHS report in which 24% of the children underweight(11),Nepal,27%(27), Vietnam, 27.7%(28), Kenya, 20%(29),Bulehora 29.2%(28),West Gojjame zone,49.2%(30), Lalibela 25.6% (31) were underweight. This variation might be due to the difference in the study area, period, and socio-economical differences of each study area.

But Research finding showed a higher prevalence of underweight than Mongolia in which the prevalence of underweight was 4.7% (32), Nigeria, 9.5% (33) and EDHS2016 report(11), which was 10%of children were underweight these may be varying according to the geographical location and the study population and health policy difference between the two countries.

As observed from this literature, there were improvements in nutrition over time. This could be attributed to the efforts of the health sector to enhance good nutritional practices through health education and the provision of micronutrients to the most vulnerable group. Also, the health extension program has included nutrition as one part of its packages. The prevalence of underweight was peaking at the age groups of 12–23 months as compared to other age groups. Our finding is consistent with a study conducted in the northern part of Ethiopia in which undernutrition was found to be peak (66.7%) at the age of 12–23 months (26). It is inconsistent with the study conducted at Butajira where underweight was reported to be peaking at 6-12 months,21.2% (26). It is also consistent with the results of researches in other African countries Congo (34)and South Africa where underweight was associated with age less than 12 months (35). This might be due to the poor habit of breast-feeding, against malnutrition.

Birth interval was independently associated with underweight as children born within 24 months of the preceding siblings were more likely to be underweight than those who born after 24 months. This study also identified that children born to a household with a low family income were more likely to be underweight as compared to children from a high family income. This finding is in line with the study conducted in Vietnam and Bangladesh (34, 36)

The children’s factors were also found to be independently associated with undernutrition among children. Male Sex was found to be significantly associated with underweight. Similarly, many studies in Ethiopia and elsewhere have reported that under-five male children are more likely to become underweight than their female counterparts (36-39). This could be because boys are more influenced by environmental stress than girls(40). The presence of diarrheal morbidity in the last two weeks before the data collection period and Diarrheal morbidity within two weeks was significantly associated underweight. The results of this study are in agreement with the results of studies conducted in different developing countries (37, 41). This is because diarrhea may result in lower appetite and poor digestion and mal-absorption.

## Conclusion and recommendation

This community-based cross-sectional study showed that the overall prevalence of underweight was relatively high among children aged six to fifty-nine months. The findings of our study showed that being male, family income below 1596 ETB, birth interval below 24 months diarrheal morbidity, were found to be significant predictors of underweight. Therefore, implementing situation based intervention by focusing on supporting housewives, treatment of Diarrheal disease, encourage ANC follows up, educate on child feeding and nutrition should be strengthened.

## Declaration

## Authors’ contributions

**LAM** contributed to generate topics, writing proposals, data collection, analyses, development of the manuscript, processed publication **YW, WSS**, and **EDA**Contributed in reviewing the proposal, assist in data collection, analyses, and critically reviewed the manuscript.

## Ethical Approval

Ethical clearance was obtained from the Institutional Review Board (IRB) of Bahir Dar University, Department of Applied human nutrition and support letter was issued from Bahir Dar University then delivered to Angolela tera district Health office and finally to the respective health institution. Besides, informed consent was obtained from the parents to confirm their willingness for participation after explaining the objective of the study. The parents were notified that they had the right to refuse or terminate at any point of the interview and their name may not be mentioned and information provided by each respondent will be kept confidential

## Availability of data and materials

Data will be available upon request from the corresponding author.

## Acknowledgments

We would like to thank study participants, data collectors, and supervisors who were involved in this study and spent their valuable time responding to our study.

## Abbreviations and acronyms

ANC: Antenatal Care
AOR: Adjusted Odds Ration
BF: Breast Feeding
CI: Confidence Interval
COR: Crude Odds Ratio
ENA: Emergency Nutritional Assessment
WAZ: weight-for -Age Z-Score
SD: Standard Deviation
SPSS: Statically Package for Social Science
UNICEF: United Nations Children’s Fund
WHO: World Health Organization

